# Reproductive state switches the valence of male urinary pheromones in female mice

**DOI:** 10.1101/2022.08.22.504866

**Authors:** Caitlin H. Miller, Tess M. Reichard, Jay Yang, Brandon Carlson-Clarke, Caleb C. Vogt, Melissa R. Warden, Michael J. Sheehan

## Abstract

Internal states shape responses to sensory stimuli. Mammalian female reproductive states are understudied considering they are one of the most regular state changes in the animal kingdom. Here we examine female house mouse preferences toward male odors across the reproductive states of estrus and late-stage pregnancy. In house mice, urine scent marks are salient social odors that convey information about the sex and identity of individuals by major urinary proteins (MUPs). Males secrete a sex-specific pheromonal protein called darcin (MUP20). Additionally, genetically diverse mice secrete unique combinations of MUPs used in individual recognition. Prior work has revealed that male odors are powerful social stimuli for female mice, yet we have a limited understanding of how the valence of such odors change across reproductive states. We discovered a valence shift among estrus and pregnant females toward novel male urine, in which estrus females exhibit preference and pregnant females show strong avoidance. This valence switch also occurs toward darcin alone, providing further support for darcin as a strong sexual signal. However, when presented with familiar male urine, the approach-avoidance response disappears, even when additional darcin is added. In contrast, when an existing identity protein (MUP11) is added to familiar male urine the approach-avoidance response is recovered. This indicates that darcin in the absence of other identity information denotes a novel male and that familiar identity information present in male urine is sufficient to modify responses to darcin. Our findings suggest that the sex and identity information encoded by MUPs are likely processed via distinct, and potentially opposing pathways, that modulate responses toward complex social odor blends. Furthermore, we identify a state-modulated shift in decision-making toward social odors and propose a neural circuit model for this flow of information. These data underscore the importance of physiological state and signal context for interpreting the meaning and importance of social odors.

## Results & Discussion

Internal state shapes the meaning and importance of sensory stimuli. For example, when you’re dehydrated a glass of water is much more satisfying than it otherwise would be. Some of the most critical yet understudied state changes are those of mammalian estrus and pregnancy. Reproductive cycles represent rare state shifts that consist of highly regular hormonal changes. These changes facilitate flexible decision-making over both short and long timescales, including adaptive responses to social environments. The value of social encounters thus varies over the course of the estrus cycle and pregnancy^1–6^. While a novel male denotes a mating opportunity for a female mouse in estrus (i.e., close to ovulation)^7,8^, a novel male intruder elicits aggression in mothers^3,9,10^. Similarly, the presence of a novel male’s odor can induce pregnancy block during early stage pregnancy in house mice, a phenomenon known as the Bruce effect^11,12^.

Social odors are a central mode of communication among mammals^13–15^. In house mice, males and females scent mark their environment with urine^7,16–20^. These urine marks convey detailed social information about the sex, competitive status, and identity of individuals^8,21–28^, which mice carefully attend to^7,8,27,29,30^. Sexually receptive female mice in estrus find male urine attractive and rewarding^22,31–34^. The component of male urine that stimulates this response is darcin (MUP20), which acts as a male sex pheromone in house mice^1,7,8^. The attractiveness of this odor wanes when females enter the quiescent phase of their estrus cycle (diestrus)^1^ and induces aggression in mothers^3^. In contrast, the response of pregnant females toward male odors is underexplored, particularly during late-stage pregnancy (i.e., the 3^rd^ trimester). The final trimester of pregnancy is intriguing because females are arguably in their least sexually receptive phase, and the physiological shifts occurring during this period may prime females for profound changes that occur upon giving birth.

Here, we examine the response of female mice in estrus and late-stage pregnancy toward male social odors (**Figure 1A–B**). The objectives of this study were to: (1) investigate the valence of male social odors across reproductive states, and (2) isolate the components of male urine that drive shifts in female responses. Specifically, we were interested in the major urinary protein (MUP) components of mouse urine which convey sex and identity information. In addition to the male pheromone darcin (MUP20), genetically diverse house mice each secrete a unique subset of identity-specific MUPs (‘central’ MUPs) that are used to recognize individuals^21,23,27,35^. We can synthesize these proteins in the lab as recombinant MUPs (rMUPs)^27,36^ and present highly controlled social odor stimuli.

**Figure 1.**
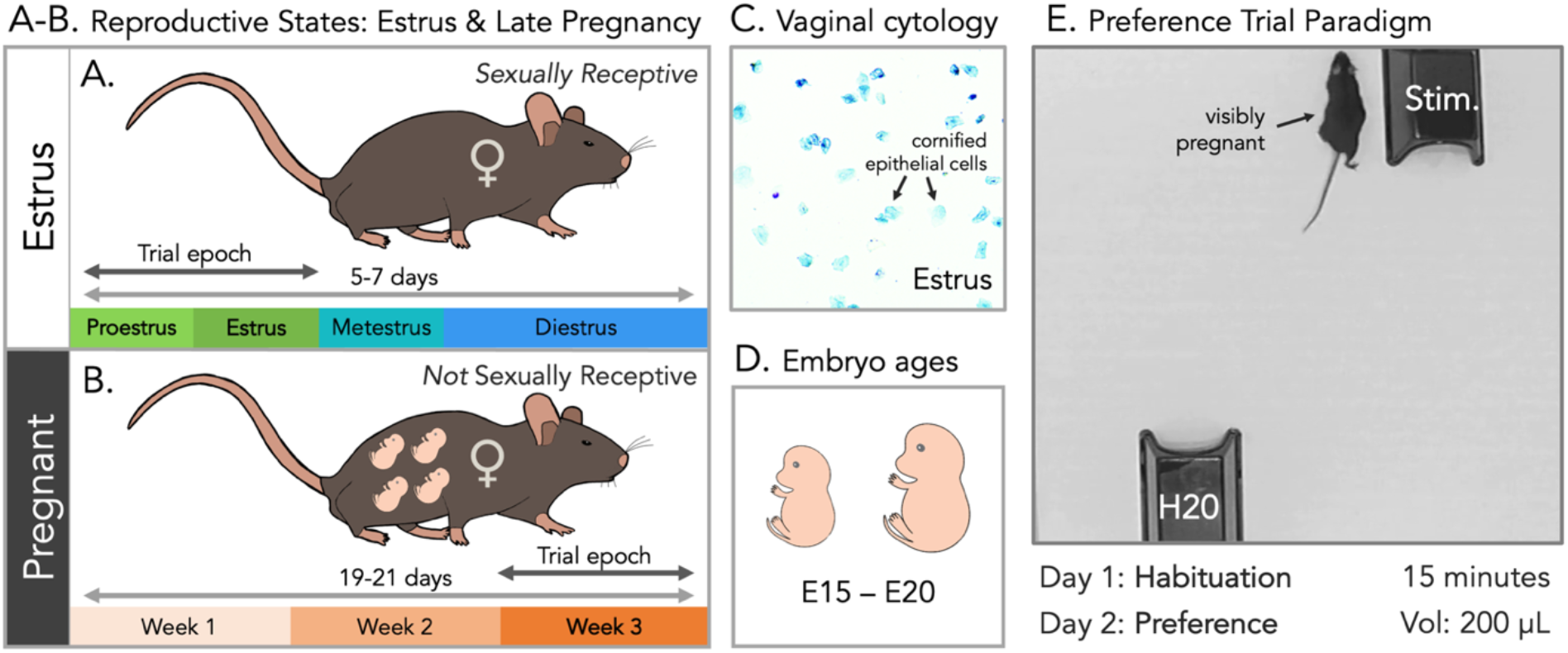
Reproductive states & experimental design. **(A)** Schematic representing the estrus cycle stages (proestrus, estrus, metestrus & diestrus) of female mice, which typically occur over the course of 5-7 days. Experimental females were close to ovulation, in the proestrus and estrus phases of the cycle, indicated with the “trial epoch” arrow (**Table S1**). **(B)** Schematic representing the 3-week duration of pregnancy. We focus on late-stage pregnancy (3rd trimester), indicated with the “trial epoch” arrow. **(C)** Vaginal cytology was used to track and stage the estrus cycles of females. Cytological swabs of females in estrus were identified by the abundance of cornified epithelial cells. **(D)** Pregnancies were staged such that we carefully tracked the progress of female pregnancies after being paired with a male and ran females with embryos at days 15 −20 of development. **(E)** Preference trial paradigm. Females were placed in an arena for 15 minutes with two huts. Under each hut 200 uL of liquid was deposited in small droplets akin to urine marking. One hut contained a social odor stimulus, and the other a water (H2O) control. Females were placed in the arena with both huts and stimuli for two consecutive days. The first day was a habituation trial, the second was a preference trial. Time spent in or on top of the huts was scored and preference indexes were calculated as the time spent in the stimulus hut relative to the water hut.

We tracked the reproductive states of naturally-cycling females using vaginal cytology to ascertain the phase of the estrus cycle^37–40^ (**Figure 1C, Table S1**). We staged and tracked pregnancies to ensure females were in their 3^rd^ trimester at the time of trials (**Figure 1D**). To examine social odor preferences, we exposed individuals to a two-day preference trial assay (**Figure 1E**). The first day was a habituation trial, as mice will explore novel environments and odors^41,42^. The second day was the preference trial, in which approach-avoidance behaviors were examined (**Table S2**). The arena held two stimulus huts: one contained a social odor, and the other contained a water control (**Figure 1E**). We calculated a preference index based on how much time females spent in or on top of the stimulus hut relative to the water control hut. A zero value indicates a female spent the same amount of time in each hut. Positive values indicate a preference for the stimulus hut while negative values indicate avoidance.

### Novel male urine and the male pheromone darcin elicit similar valence shifts across reproductive states

We first tested the response of estrus and pregnant females toward novel male urine. As all experimental subjects were C57BL/6J (C57) females, novel male urine was collected^43^ from a genetically distinct inbred mouse line (NY3)^44,45^. This was necessary to ensure that the novel male urine stimulus contained a different MUP profile from C57 males, because all individuals of a given inbred strain share identical MUP profiles^46,47^. In the wild different males secrete distinct urine profiles used to recognize individuals^21,23,27,35^, and we wanted to present ecologically relevant novel male social odors similar to prior studies^8,48^. Furthermore, all females had prior exposure to adult male C57 urine (via their father and/or stud male).

We predicted that novel male urine would be a potent social odor stimulus for females, given prior research on females in estrus^1,22,31^ and mothers^3,9,10^. Similar to what has been previously described^8,22,48^, we observed a preference among sexually receptive estrus females toward novel male urine (*t*1,14 = 2.1, p = 0.03; **Figure 2**). In contrast, late-stage pregnant females showed a strong avoidance to the same stimulus (*t*1,15 = −3.6, p = 0.001; **Figure 2**). Estrus and pregnant females also differed in their preference for novel male urine (M1: *t*1,125 = 5.1, p < 0.0001; **Figure 2 & Table S3**). Females therefore exhibit state-dependent changes in decision-making toward social information.

**Figure 2.**
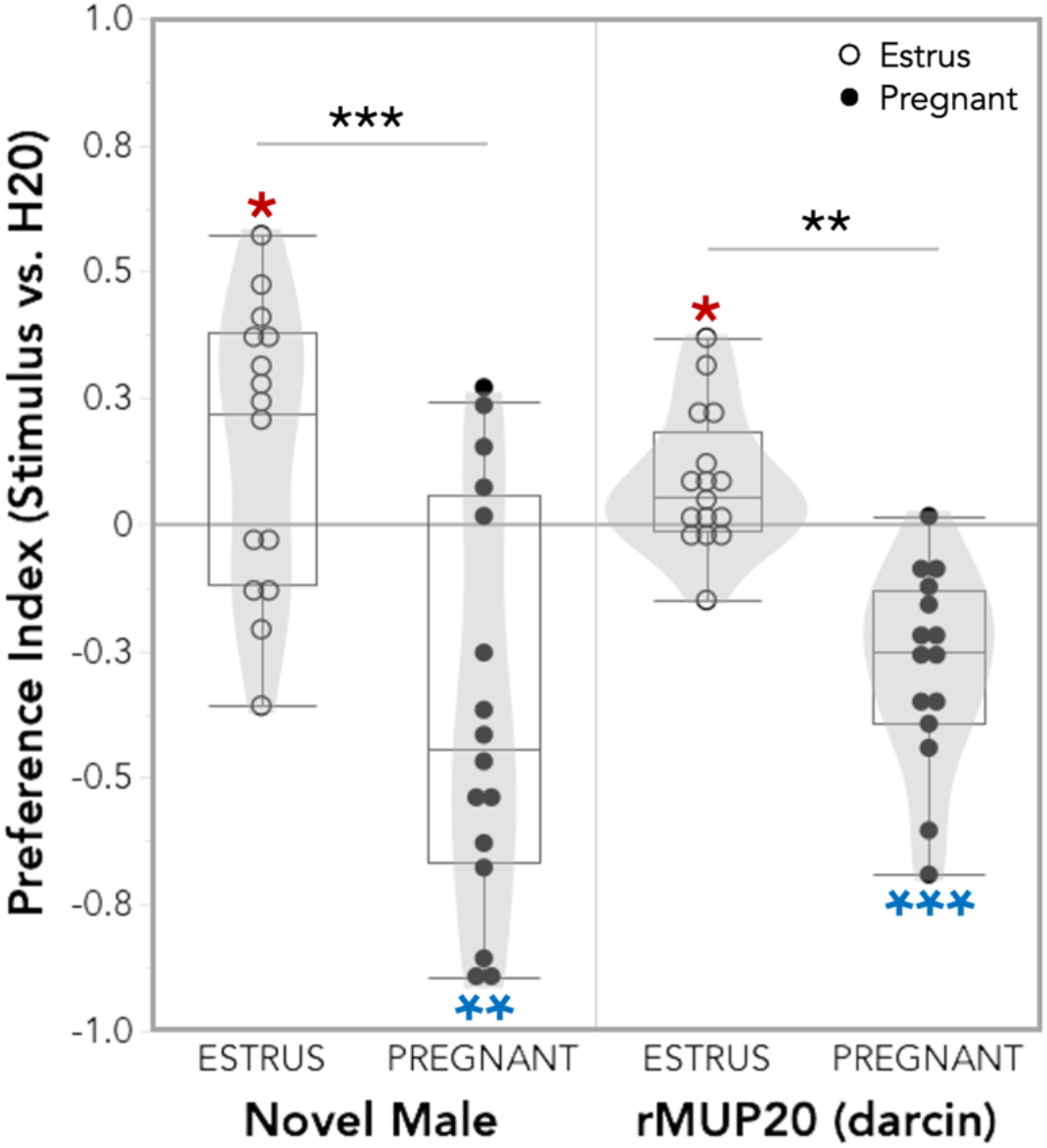
Estrus females show a preference for novel male urine and to rMUP20 (darcin), while pregnant females exhibit strong avoidance to both. All data is from the preference trial day. The preference index: how much time females spent in the stimulus hut relative to the control water hut. Stimulus huts contained either whole novel male urine or the recombinant male urinary protein rMUP20 (i.e., darcin). A zero value means a female spent the same amount of time in each hut. Positive values indicate preference for the stimulus hut, and negative values indicate avoidance. One-sample t-tests (deviation from 0) significance values are indicated with red asterisks (positive deviation or blue asterisks (negative deviation). A linear model was used to examine relationships, analyses of variance were used to test for overall effects, and pairwise comparisons were performed using the *emmeans* package. Estrus: novel male, n=15; rMUP20, n=16. Pregnant: novel male, n=16; rMUP20, n=15. Significance codes: * p<0.05, ** p<0.01, *** p<0.001.

A key component of male house mouse urine is the major urinary protein and sex pheromone darcin (MUP20). We next examined if darcin on its own was sufficient to elicit the observed approach-avoidance switch in estrus and pregnant females. Following previous studies^1,27^, we synthesized and presented recombinant darcin (rMUP20) as a stimulus. This single urinary protein elicited a similar valence shift across female states when compared to novel male urine (**Figure 2**). We found that females in estrus were attracted to darcin, as previously reported^1,8,48^ (*t*1,15 = 2.5, p = 0.01; **Figure 2**). In contrast, pregnant females displayed robust avoidance (*t*1,14 = −5.7, p = 3e-05; **Figure 2**). Females also differed in their preference for darcin across reproductive states (M1: *t*1,125 = 3.6, p = 0.009; **Figure 2 & Table S3**). This response provides further evidence that darcin is a strong social signal in house mice and is a clear example of a valence switch toward a specific and salient sex pheromone.

### Altering the identity profile of familiar male urine recovers the approach-avoidance valence switch observed toward novel male urine

There is, however, an important caveat to these results which required further probing – adult males typically secrete darcin in their urine^23,24,49^. This includes the pregnant female’s stud male that was co-housed with her during trials. Therefore, darcin itself is unlikely to be aversive to pregnant females because they are consistently exposed to darcin by their stud male. We hypothesized that the observed response toward darcin alone may be due to the absence of other key identity information typically associated with their stud male’s urine (i.e., his genotype-specific identity MUP expression pattern).

To address this, we presented pregnant females with urine from their C57 stud male, and estrus females with adult C57 male urine. C57 male urine is not novel to virgin estrus females as they were exposed to it via their fathers. Moreover, because they are from the same inbred line, C57 males and females produce nearly identical MUPs^46,47,49^. The similarity in urine profile conveys relatedness, and females will preferentially mate with unrelated males based on their MUP profile^50^. We also manipulated the MUP content of stud/C57 male urine by adding equivalent amounts of either the male-specific recombinant protein darcin (rMUP20) or a recombinant identity MUP present in C57 urine (rMUP11)^27,49^. In doing so, we effectively doubled the amount of MUP20 or MUP11 present in the C57 male urine. We predicted that adding rMUP20 to familiar C57 male urine would not alter the urine identity profile, and therefore would not elicit approach-avoidance behaviors in females. Conversely, as the relative ratios of identity MUPs are to recognize individuals^27^, we predicted that adding more rMUP11 would alter the identity of the male urine and thus provoke a valence switch. Importantly, in both cases we added peptides already present in the urine stimuli, and thus only changing the ratios of MUPs.

When presented with unaltered stud/C57 male urine, neither estrus (*t*1,16 = 0.21, p = 0.83) nor pregnant females (*t*1,16 = 1.1, p = 0.31) displayed any preference, and females did not differ across states (M1: *t*1,125 = −0.90, p = 0.99; **Figure 3 & Table S3**). For estrus females, this is likely because C57 male urine smells like that of a full sibling^50^, and is a less attractive mating opportunity in comparison to the urine of a male with a different genotype. For pregnant females this is a male they see daily in their home cage, and hence is a highly familiar odor. The response to stud/C57 male urine with added rMUP20 also revealed no differences across reproductive states (M1: *t*1,125 = −0.14, p = 1.0, **Table S3**), and no approach or avoidance among estrus (*t*1,9 = 0.93, p = 0.38) or pregnant females (*t*1,10 = 1.6, p = 0.14; **Figure 3**). This shows that adding additional darcin to familiar male urine does not alter the identity or attractiveness of the stimulus for females. In contrast, when rMUP11 is added to familiar male urine the response toward novel male urine is recovered, and females significantly differ in preference across reproductive states (M1: *t*1,125 = 3.1, p = 0.04; **Figure 3 & Table S3**). Estrus females displayed approach behavior (*t*1,8 = 2.0, p = 0.039), and pregnant females exhibited avoidance (*t*1,8 = −1.9, p = 0.048; **Figure 3**). It is notable that shifting the ratio of a single identity MUP (MUP11) was sufficient to elicit this response.

**Figure 3.**
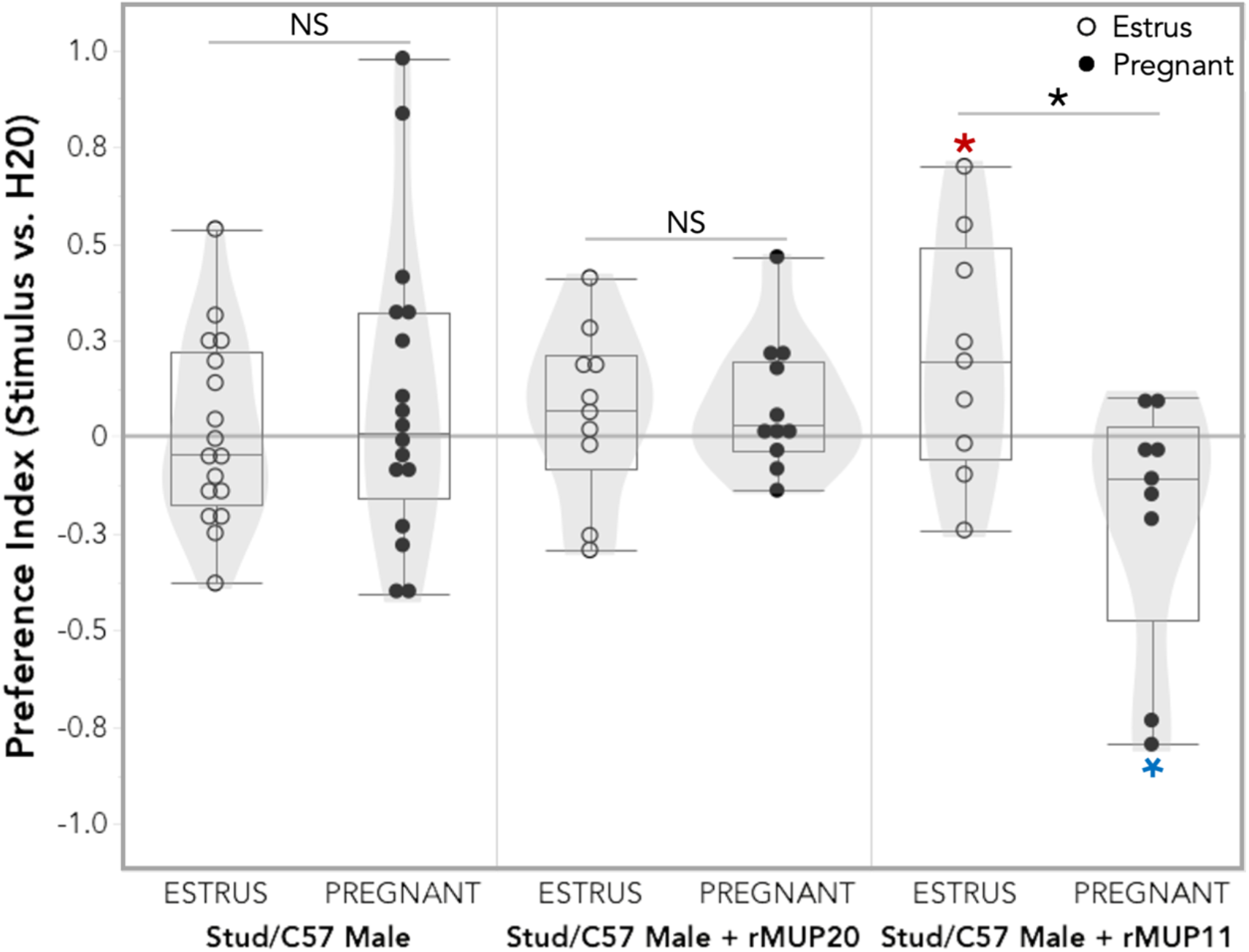
Approach-avoidance behaviors of estrus and pregnant females toward whole stud/C57 male urine, or stud/C57 male urine manipulated with added darcin (rMUP20) or rMUP11. The preference index: how much time females spent in the stimulus hut relative to the control water hut. A zero value means a female spent the same amount of time in each hut. Positive values indicate preference for the stimulus hut, and negative values indicate avoidance. One-sample t-tests (deviation from 0) significance values are indicated with red asterisks (positive deviation or blue asterisks (negative deviation). A linear model was used to examine relationships, analyses of variance were used to test for overall effects, and pairwise comparisons were performed using the *emmeans* package. Estrus: C57, n=17; C57+rMUP20, n=10; C57+rMUP11, n=9. Pregnant: Stud, n=17; Stud+rMUP20, n=11; Stud+rMUP11, n=9. Significance codes: NS p>0.05, * p<0.05.

### Sex and identity information are processed distinctly: a proposed model

Our results reveal that darcin on its own denotes the presence of a novel male. This appears to be driven by the absence of identity information associated with a familiar male. This suggests that sex and identity information are processed via distinct, and potentially opposing, sensory pathways that converge to modulate behavioral responses. Given the observed preference patterns and existing circuit knowledge, we propose a model of information flow (**Figure 4**).

**Figure 4.**
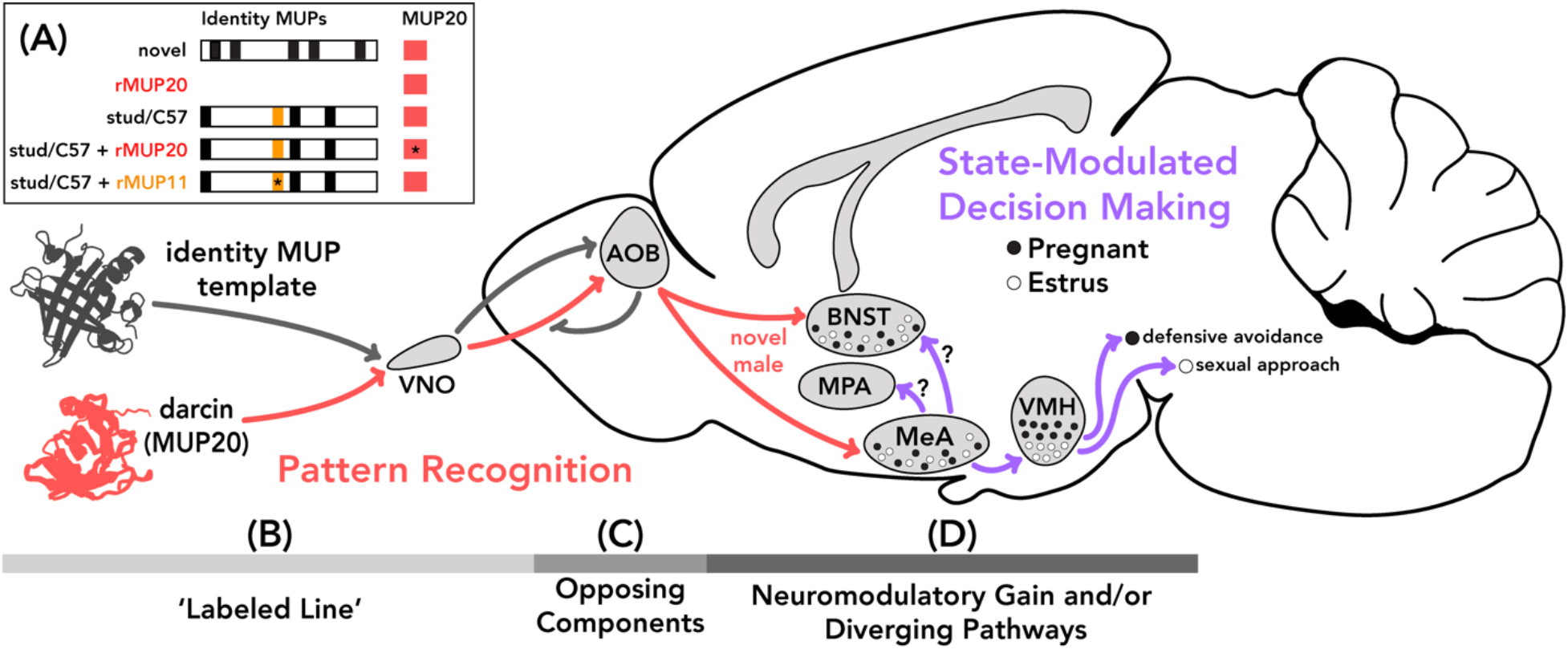
Model of information flow mediating pattern recognition and state-modulated decision-making in female mice toward social odors. **(A)** Schematic depicting the MUP components of male urine and the recombinant MUP stimuli. For the ‘central’ identity MUPs protein expression is indicated by black bars, except for MUP11 which is indicated with a yellow bar. Novel male urine has a distinct MUP array compared to familiar stud/C57 male urine. A red bar indicates the presence of the male-specific pheromone MUP20 (darcin). All male urine contains darcin. For manipulated urine stimuli containing additional rMUP an asterisk indicates the added rMUP (rMUP11 or rMUP20). **(B)** Model for the distinct processing of sex (red) and identity information encoded by MUPs (black), in which the male pheromone darcin (red) has specialized circuitry (‘labeled line’) projecting from the peripheral olfactory vomeronasal organ (VNO) to the accessory olfactory bulb (AOB). **(C)** Templates for familiar male identities are stored in the AOB. When the MUP identity matches the olfactory memory template, this initiates a suppression of darcin-specific signaling cascade. However, if this identity pattern recognition does not occur, the darcin signaling cascade continues to downstream limbic regions and conveys the presence of a novel male. **(D)** Female responses to novel male urine are modulated by their hormonal reproductive state. Females in estrus show a preference for novel male urine, and pregnant females exhibit avoidance. This approach-avoidance valence switch toward the same odor stimuli is likely modulated by activity in one or both of the primary projection targets downstream of the AOB: bed nucleus of the stria terminalis (BNST) or the medial amygdala (MeA). State modulated decision-making is likely mediated by shifts in neuromodulatory gain toward changing hormone levels within the MeA. The MeA in turn gates information to distinct subregions of the ventromedial hypothalamus (VMH) via diverging projections, such that distinct VMH populations are activated during sexual approach (estrus) as opposed to defensive avoidance (pregnant). Alternatively, information may be gated to different downstream limbic brain regions such as the medial preoptic area (MPA) and/or the BNST.

The robust responses observed toward darcin in this study and by other groups^1,7,8,48^, suggest a specialized circuitry (‘labeled-line’; **Figure 4B**). This is likely given that a related pheromone (ESP1) which stimulates sexual posturing in females is detected via a labeled-line^51,52^. Our results demonstrate that this proposed darcin-specific signal cascade is interrupted if a familiar identity-matched ‘central’ MUP template is present. Darcin is therefore not part of the identity template, but instead acts as a robust male signal that gets suppressed by the presence of familiar male identity information.

Our results provide further evidence that individual recognition is determined by combinatorial coding of ‘central’ identity MUPs^27,53^, as the manipulation of a familiar male MUP ratios with a single identity rMUP (rMUP11) stimulated a response similar to that of a novel male (**Figure 4A**). In this manipulation the template did not match (due to the altered ratios of identity signaling peptides) and thus the darcin signal cascade was not suppressed. Signal context is consequently crucial for mediating adaptive social and spatial preferences, particularly when determining the presence of a male in the environment and whether or not that male is familiar.

Pheromonal suppression via individual pattern recognition likely occurs in the accessory olfactory bulb (AOB; **Figure 4**), as MUPs are detected by the accessory olfactory (i.e. vomeronasal) system^1,27,48^ and stud male odor memory formation requires a functional AOB but not other downstream limbic regions^12,54,55^. Evidence suggests that this sensory memory is maintained by local modulation of AOB projection neuron membrane excitability^56–58^. We propose a local inhibition of the darcin ‘labeled-line’ within the AOB (i.e., opposing components; **Figure 4C**), such that familiar male identities block the darcin circuit cascade, and only upon detection of a novel male are strong neuronal signals sent downstream.

The state-modulated valence switch in decision-making toward novel male odors in estrus and pregnant females likely occurs in the primary downstream limbic targets of the AOB: the medial amygdala (MeA) and/or the bed nucleus of the stria terminalis (BNST). The MeA and BNST receive abundant direct excitatory input from the AOB and have hormone-sensitive neuronal populations^59–62^. These brain regions are primed to facilitate shifts in responses to varying hormone levels by neuromodulatory gain and/or diverging pathways.

The MeA is a likely target for such modulation given its central location within limbic circuitry, and its role in gating responses to reproductive and defensive stimuli^63,64^. Furthermore, the MeA is strongly activated by unfamiliar relative to familiar male urine in female mice^65^, and specific MeA neuronal populations are activated by darcin^7^ and ESP1^51^. The MeA in turn sends prominent projections to the ventromedial hypothalamus (VMH), which has abundant estrogen and progesterone-sensitive neurons^51,66–78^. In females, two distinct VMH subpopulations are differentially activated by mating^68,78^ and fighting^68^, reminiscent of the approach-avoidance behaviors observed across reproductive states. Moreover, a recent study identified a VMH subpopulation in female mice that modulates sexual receptivity in an activity-dependent manner^78^. We propose that neuromodulatory gain within the MeA gates input to the VMH, and thereby differentially activates distinct subpopulations in a state-dependent manner (**Figure 4D**). Alternatively, information may be gated to other brain regions. For example, MeA projections to the BNST may mediate estrus female approach^79^, and projections to the VMH may mediate pregnant female avoidance^71,79,80^.

This preference paradigm and circuit model provide a unique opportunity to interrogate the neurobiological mechanisms underlying valence processing and hormonal state shifts. The behavioral and physiological changes that occur over the course of reproductive cycles are uniquely regular. Furthermore, pregnancy is one of the most dramatic physiological changes that can occur within an individual’s lifetime, and yet is sorely understudied. These results highlight the biomedical importance for considering reproductive state variation as well as how social odor assays are utilized and interpreted. Our findings reveal key insights into how sex and identity information are processed, and open new avenues for investigating recognition mechanisms.

## Material & Methods

### KEY RESOURCES TABLE

| REAGENT or RESOURCE | SOURCE | IDENTIFIER |
| --- | --- | --- |
| *Experimental Models: Organisms/Strains* |  |  |
| C57BL/6J (*M. m. domesticus*) | The Jackson Laboratory | JAX: 000664 |
| *Recording & Imaging Systems* |  |  |
| iDVR-PRO CMS IR security cameras | CCTV Camera Pros | https://www.cctvcamerapros.com/DVR-Remote-Access-non-IE-s/470.htm |
| 40X-2500X Plan Infinity Kohler Laboratory Trinocular Compound Microscope + 18MP USB3.0 Camera | AmScope | T720QC-18M3 |
| *Materials & Solutions* |  |  |
| Whatman filter paper (46 x 57 cm) | Sigma-Aldrich | 1002-917 |
| Gibco Phosphate Buffered Saline pH 7.4 (1X) | Thermo Fisher Scientific | 10010031 |
| JorVet Dip Quick Stain Kit | Jorgensen Labs | J0322 |
| pMAL Protein Fusion and Purification System | New England Biolabs | E8200 |
| *Software and Algorithms* |  |  |
| BORIS 8.0 | Behavioral Observation Research Interactive Software | https://boris.readthedocs.io/en/latest/ |
| R v.3.6.3 | The R Project for Statistical Computing | https://www.r-project.org/ |
| R package *trajr* | 81 | https://cran.r-project.org/web/packages/trajr/index.html |
| R package *lme4* | 82 | https://cran.r-project.org/package=lme4 |
| R package *lmerTest* | 83 | https://CRAN.R-project.org/package=lmerTest |

### EXPERIMENTAL MODEL AND SUBJECT DETAILS

#### Animals

All experimental subjects in this study were C57BL/6J female mice (n=135) obtained from The Jackson Laboratory (JAX stock #000664). At the time of experimentation all females were fully adult (2-5 months old), estrus females were virgin, and pregnant females were in the third week of their first pregnancy. Virgin females used to track estrus cycles were housed in holding cages with other females. Pregnant females were housed in breeding cages with their stud C57BL/6J (C57) male. All holding and breeding cages contained corn cob bedding, cardboard huts, and cotton nestlets. Mice were housed in an Animal Care facility at Cornell University with a 14:10 shifted light:dark cycle (dark cycle: 10AM–8PM), with food and water provided *ad libitum.* To reduce handling stress confounds, mice were transferred between their home cage and the experimental arena using transfer cups^84^.

### METHOD DETAILS

#### Stimuli

The social odors used as trial stimuli consisted of whole urine and/or recombinant major urinary proteins (rMUPs) synthesized in lab: novel male urine, stud/C57 male urine, darcin (rMUP20), stud/C57 male urine with added rMUP20, and stud/C57 male urine with added rMUP11. While darcin is a male-specific major urinary protein^23,24,85^, mice secrete many other urinary proteins that mediate identity recognition (also called ‘central’ MUPs)^21,23,24,27,35^. In the wild, individuals within a population typically excrete distinct combinations of MUPs^23,86^. Individuals within an inbred strain secrete the same combination (as a result of genetic inbreeding)^46,47^. C57 mice typically express a MUP with molecular weight 18693 Da matching the sequence of MUP11^27,49^. Accordingly, rMUP11 was used to shift the identity protein profile of C57 male urine by changing the ratio of rMUP11 present in the familiar C57 male urine stimulus. For stimuli with added rMUPs we effectively doubled the amount of the ‘manipulated’ MUP present in the urine (thereby altering the MUP ratio). Stimuli were all presented as a total 200 μL volume that was aliquoted in small 25-50 μL droplets to mimic urine marking under specific stimulus huts in the preference trial arena (**Figure 1E**).

#### Urine collection

Urine collection was performed using the single animal method: males were placed atop a metal grate (an upside down cage hopper) over a clear plastic bag for 30 minutes to 1 hour^43^). Mice were subsequently taken off the plastic bag and returned to their home cage. The urine droplets present on the plastic bag were collected and stored at −80°C until use. Two types of urine were collected: novel male urine and stud or C57BL/6J male urine. Novel urine was collected from a distinct inbred mouse line (NY3: this strain is related to the SarA/NachJ, SarB/NachJ and SarC/NachJ strains available from the Jackson Lab^44^), to ensure that the urinary protein profile was completely novel to experimental females^21,23,27,46^. For novel male urine we collected a large batch of urine from over 20 adult individuals for each stimulus. This was done such that the exact same novel male genotype urine stimulus was presented to experimental females across trials, without any individual-specific effects of urine odors. Stud and pooled C57BL/6J male urine was also collected. For stud male urine, we collected urine from a pregnant female’s specific individual C57BL/6J stud male a week prior to the start of the two-day preference trial. Virgin estrus females by definition did not have a stud male. They were presented adult C57BL/6J male urine. We collected a large batch of urine from over 20 adult male C57BL/6J genotype mice, again to prevent any individual-specific effects of urine odors such that all estrus females received the same C57BL/6J male urine stimulus. For all urine collection, urine was stored on the day of collection at −80°C. Once a sufficient volume was collected to use as stimuli, individual aliquots were thawed on ice, and urine was pooled into a single volume and subsequently aliquoted into trial appropriate volumes (200 μL) and stored at −80°C until use.

#### Major urinary protein synthesis

Using complementary DNA (cDNA) libraries from adult male house mice, complete MUP cDNAs were obtained from liver tissue. MUP cDNAs were amplified and sequences were checked using Sanger sequencing methods.These MUPs were cloned into an *E. coli* C2523 pMAL-c5X vector and recombinant MUP (rMUP) was made using pMAL Protein Fusion and Purification System (New England Biolabs) using methods similar to prior studies^27,36^. Mature MUP11 (ENSMUSP00000095654.4) and MUP20 (ENSMUSP00000073667.4) lacking signal peptide^49^ were produced as rMUPs to be used as stimuli in preference trials^27,36^. To ensure rMUPs were successfully made, an SDS-PAGE gel was run to verify protein length and compared to male mouse urine. When rMUPs were added to stimuli in behavioral trials, they were presented at a concentration of 5 mg/ml which is comparable to typical urinary protein concentrations found in mouse urine^27^.

#### Vaginal cytology

Rather than removing the ovaries of females and performing hormonal treatments, we wanted to track the natural estrus cycles of females to observe the natural hormonal changes that occur in female mice. Vaginal cytology was used track the estrus cycles of females using standard procedures^37–40^, and was performed 1-2 hours prior to the start of the dark cycle (8-9 AM). Vaginal cytology was performed daily for at least two full estrus cycles for each female to ensure careful timing of the proestrus-estrus phase during preference trials. Mice were handled gently to reduce stress, restrained only by the base of their tail and were allowed to rest their forepaws on the cage hopper. A pipette was filled with 25 μL of Gibco Phosphate Buffered Saline (PBS) pH 7.4 (1X), and then gently lavaged at the vaginal opening. The pipette was flushed approximately ten times, or until the solution was cloudy, indicating the presence of cells. The solution was carefully released and spread onto a glass slide. Each mouse had its own slide and pipette to avoid sample contamination. The glass slides were stained with JorVet DipQuick Stain Fixative, Stain Solution, and Counter Stain. Extra stain was rinsed off with a water dip, and subsequently examined under a 10X resolution microscope (AmScope: T720QC-18M3). Each stage of the estrus cycle is dictated by which cell types are present, leukocytes, cornified epithelial cells, and nucleated epithelial cells, and the proportion of cell types present^37–40^ (**Table S1**). Specifically, we looked for females with cornified epithelial cells or nucleated epithelial cells, indicating they were in estrus^37–40^ (**Table S1**). As the trials were two days long, on the first habituation trial day we used females that were in the earlier stages of estrus (proestrus or their first day of estrus). On the second preference trial day we timed it such that females were in peak estrus (and close to ovulation).

#### Pregnancy staging

We set up timed pregnancies because we were interested in studying late stage pregnancy in female mice (females in their 3^rd^ and final week of gestation: 15-21 days pregnant^87^). To do this, adult virgin female mice (at least 1.5-2 months old) were paired with an adult male mouse on a specific date, and their subsequent date for giving birth was predicted to be approximately three weeks after initial breeding cage setup (gestation time of 19-21 days). The day after breeding cages were setup, females were checked to see if she showed signs of successful mating indicated by the presence of a copulatory plug^88^. Females did not always show a plug, so their size was monitored daily until the day of birth^89^. In particular, we carefully tracked the date in which females became visibly pregnant, as this was very predictive of being in their second week of pregnancy. The combination of the predicted birth date based on the breeding cage set up, along with the tracked pregnancy bulge, provided a reliable guide to the pregnancy stage of females. When females were sufficiently large and between 15 and 21 days after the being paired with a male, females entered into the two-day preference trial assay. All pregnant females successfully gave birth, and the date of birth was used to accurately determine the stage of pregnancy and embryo development (E1-E20) at the time of experimentation.

Females were excluded from trials if they were no longer in estrus on their preference trial day, if pregnant females gave birth on their preference trial day, or if females were actually in their 2^nd^ trimester during the trial period (as determined by the date of birth of their pups).

#### Preference trials

Because house mice are primarily nocturnal^90^, trials occurred during their active dark cycle period, and all experimentation occurred between 11 AM - 2 PM to minimize circadian variation. All trials were performed in two trial chambers that were sound-proofed and fitted with security camera recording systems (iDVR-PRO CMS; 1080p; 30 fps). PVC Trial arenas (50 cm x 50 cm) were placed into these sound-proofed chambers. Large sheets of Whatman filter paper lined the floor of each trial to control the placement and presentation of social odor stimuli. The arena contained two opaque red plastic huts, placed in the lower left corner and the upper right corner. Underneath each hut 200 uL of stimulus or a control were pipetted onto the filter paper in small 25-50 uL droplets, to mimic the pattern of a mouse urine scent marking. In each trial, one hut contained a social odor stimulus (e.g. novel male urine) and the other contained water as a control. The location of the stimulus odor was randomized. Stimuli were aliquoted just prior to the start of the trial. Trials were performed in under mildly aversive full spectrum light to encourage investigation of the huts.

The preference trial assay consisted of two days, in which the mouse was exposed to the same arena and the same stimuli (in the same locations) on both days. The first day was a habituation trial, as mice will investigate novel environments and odors. The second day was the preference trial, for which approach-avoidance behaviors were examined. Once a mouse was trial ready (i.e., females in late pregnancy or in estrus), mice were transferred in a cup from their holding cage to the center of the arena, to reduce stressful handling. All trials were 15 minutes in duration. At the end of the trial, females were gently removed from the trial arena and returned to their home cage. Between trials, arenas and all chamber surfaces were cleaned thoroughly with 70% EtOH. Experimental females were used once for one preference trial stimulus trial series (i.e., no females were reused across reproductive states or stimuli).

#### Behavioral scoring and analysis

Behavioral trials were scored using Behavioral Observation Research Interactive Software (BORIS)^91^. Trials were scored blind to the stimulus presentation. The mouse’s movements were observed and classified as either under or above the huts (state events; **Table S2**). These classifications were further distinguished on the basis of whether they occurred in the front or back hut in the arena, which contained different stimuli (social stimuli or water control). Using the scored spatial data, similar to a prior study^1^ preferences indices (PIs) were calculated as follows: PI = (Σ Time spent investigating stimulus - Σ Time spent investigating control) / (Σ Time spent investigating stimulus + Σ Time spent investigating control). Investigations include time spent underneath or on top of huts. Zero value means they spent the same amount of time in each hut. Positive values indicate preference for the stimulus hut (more time was spent in the stimulus hut), and negative values indicate avoidance for the stimulus hut (more time was spent in the water hut).

### QUANTIFICATION AND STATISTICAL ANALYSIS

We conducted all statistical analyses in R 3.6.0 (R Development Core Team 2019). We used linear modeling and paired statistical tests to examine relationships between dependent variables (reproductive state and social odor stimuli) and response variables (preference indices). Models were fitted using the package *lme4*^82^. The *lmerTest* package was used to calculate degrees of freedom (Satterthwaite’s method) and p-values^83^. We used a type 3 analysis of variance (ANOVA) to test for overall effects of fixed factors or interactions in the models. Post hoc comparisons were conducted using the *emmeans* package^92^. One-sample t-tests were performed to examine whether experimental groups preference indices deviated significantly above or below zero. R script and data sheets used for all statistical analyses are provided.

## Declarations

### Ethical statement

All experimental protocols conducted at Cornell University were approved by the Institutional Animal Care and Use Committee (IACUC: Protocol #2015-0060) and were in compliance with the NIH Guide for Care and Use of Animals.

### Data accessibility

Data sheets and R code used in all analyses are available on the Dryad Digital Repository.

### Author contributions

CHM and MJS conceived the study. CHM performed trials and analyses. TMR, JY and BCC tracked reproductive cycles and scored behavioral trials. CCV developed code for video tracking. CHM wrote the initial drafts of the paper. MJS and MRW edited the manuscript. All authors contributed to manuscript preparation.

## Competing interests

The authors declare no competing interests.

## Funding

This research was funded by USDA Hatch Grant (NYC-191428; Michael Sheehan) The funders were not involved in the design of the study; the collection, analysis and interpretation of data; the writing of the manuscript and any decision concerning the publication of the paper.

## Acknowledgements

We thank Kevin Besler, Christen Rivera-Erick, Melanie Colvin and Jeremy Cusker for crucial technical assistance; Russell Ligon for helping establish recording systems and tracking methods in the lab; Eileen Troconis for helpful manuscript feedback.

## Supplemental Data

**Table S1.**
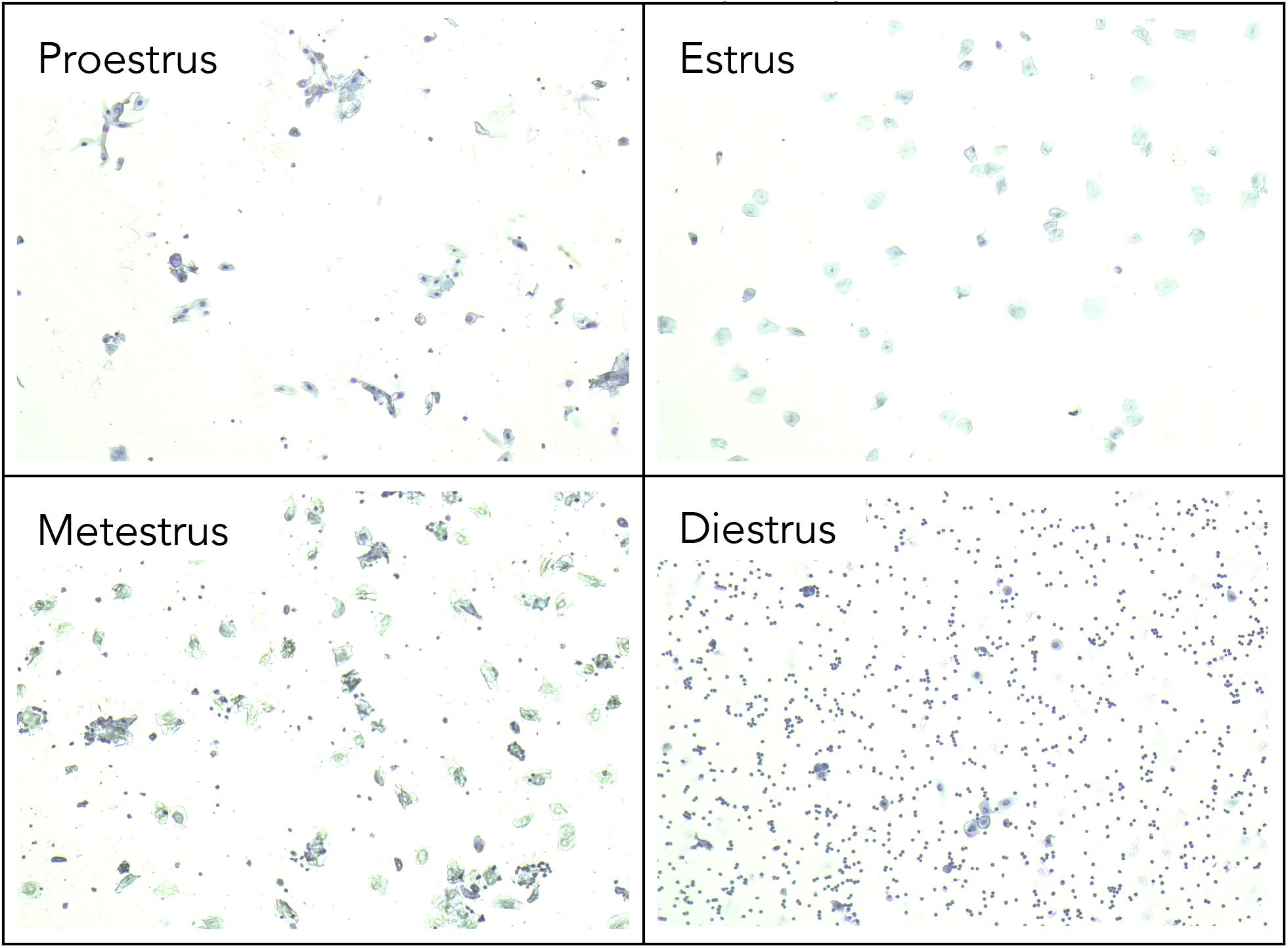
Estrus cycle vaginal cytology images. Vaginal swabs of females in proestrus are characterized by nucleated epithelial cells. Females in estrus have primarily cornified epithelial cells. As females enter the quiescent phase there are more leukocytes, as observed in metestrus. Females in diestrus have almost exclusively leukocytes.

**Table S2.** Behavioral ethogram for scoring preference trials.

| Behavior | Description |
| --- | --- |
| Under Hut | Mouse goes under the hut and stays under it for a period of time. Behavior was modified based on which hut the mouse went under—front or back hut. |
| Above Hut | Mouse climbs on top of the hut and remains there for a period of time. Behavior was modified based on which hut the mouse was above—front or back hut. |

**Table S3.** Analysis of variance (ANOVA) table for the linear model (M1) used to model female preference across reproductive states and stimuli. The response variable is the preference index (PI). Significance codes: * p<0.05, ** p<0.01, *** p<0.001.

|  | **M1 : Preference Index ~ State \* Stimulus** |  |  |  |  |
| --- | --- | --- | --- | --- | --- |
|  | Df | Sum Sq | Mean Sq | F Value | Pr (>F) |
| State | 1 | 1.8275 | 1.82748 | 22.8322 | 4.869e-06 *** |
| Stimulus | 4 | 0.8532 | 0.21329 | 2.6648 | 0.03552 * |
| State : Stimulus | 4 | 2.1105 | 0.52763 | 6.5921 | 7.582e-05 *** |
| Residuals | 125 | 10.0049 | 0.08004 |  |  |

## Notes

### Competing Interest Statement

The authors have declared no competing interest.

